# Harnessing Human Connectome Project Data for Replication Studies: A Demonstration for Resting State-Task Network Correspondence

**DOI:** 10.1101/111021

**Authors:** Lisa D. Nickerson

## Abstract

After a series of reports uncovered various methodological problems with functional magnetic resonance imaging (fMRI) research, considerable attention has been given to principles and practices to improve reproducibility of neuroimaging findings, including promotion of openness, transparency, and data sharing. However, much less attention has been given to use of open access neuroimaging datasets to conduct replication studies. A major barrier to reproducing neuroimaging studies is their high cost, in money and labor, and utilizing such datasets is an obvious solution for breaking down this barrier. The Human Connectome Project (HCP) is an open access dataset consisting of extensive behavioral and neuroimaging data from over 1,100 individuals and there are numerous ongoing HCP-harmonized studies of lifespan and disease that will ultimately release data through HCP infrastructure. To bring attention to the HCP and related projects as an important resource for conducting replication studies, I used the HCP to conduct a replication of a highly cited neuroimaging study that showed correspondence between resting state and task brain networks.

---

Recent reports on reproducibility in psychology research have produced alarming findings that have drawn attention to this issue in many scientific disciplines, not just psychology.^1,2,3^ The Open Science Collaboration (2015) attempted to replicate 100 psychology studies but succeeded in replicating only 39,^4^ and a survey by the journal Nature of 1,576 researchers showed that more than 70% of researchers have tried and failed to reproduce another scientist’s experiment and more than half reported failing to reproduce their own experiments.^5^ Various strategies have been proposed to improve the reliability and efficiency of scientific research^2^, including the use of registered reports, adopted by more than 40 journals to date to both enhance and incentivize reproducibility.^6^ The field of neuroimaging using functional magnetic resonance imaging (fMRI) has come under fire many times in the past decade when research meant to evaluate best practices in data analysis turned up several dramatic problems. For example, work by Vul et al. (2009) found that extremely high correlations between brain activation and personality measures in a large collection of neuroimaging studies arose because of circular analysis practices,^7^ work by Bennett et al. (2010) reported brain activation during a social cognition task in a dead salmon as an illustration of the inadequacy of commonly used corrections for multiple comparisons,^8^ and a recent publication by Eklund et al. (2016) revealed that parametric statistical methods implemented in many common neuroimage analysis software packages are invalid for cluster-wise inference, calling into question findings from a number of fMRI experiments.^9^ This last report has generated a great deal of controversy^10–14^ (see also http://www.ohbmbrainmappingblog.com/blog/keep-calm-and-scan-on) and has drawn attention to efforts within the neuroimaging community to identify best practices for fMRI research, from study design to data collection, to data analysis (http://www.humanbrainmapping.org/cobidas).

Most recently, Nature Neuroscience published a Focus issue on Human Brain Mapping that presents several articles discussing these issues in more detail, including how to define what constitutes reproducibility in fMRI research.^15,16^

While considerable attention is being given to the principles and practices to improve reproducibility in neuroimaging studies using MRI, including data sharing^17^ and existing repositories (see NeuroImage issue 1:24 (Part B) for detailed descriptions of several repositories^18^), less attention is being given to the actual existing neuroimaging datasets that are now available to facilitate such studies.^19–23^ Indeed, a major barrier to conducting reproducibility studies is the cost in both money and labor that it takes to perform neuroimaging studies, and these open datasets represent one of the best options for breaking down this barrier, in addition to presenting opportunities to publish original research. A recent study was published that perfectly illustrates the power of open access data and how we, as researchers, can think about using shared data to replicate our research, while still conducting original research. Zhao et al. (2017)^24^ drew data from one existing neuroimaging repository^23^ to conduct an original research study on the relationship between cortical thickness and neuroticism in individuals with alcohol use disorder, then drew data from a different neuroimaging repository^22^ to replicate the findings. Such studies demonstrate how replication truly enhances any research study by making publications more compelling and increasing our confidence in the findings.

One of the datasets used in the aforementioned study by Zhao et al (2017) is the data from the Washington University-University of Minnesota (Wu-Min) Human Connectome Project (HCP),^22^ an NIH-funded initiative to map the human brain connectome. In brief, behavioral and neuroimaging data were collected in more than 1,100 participants and the HCP have made the data publicly available (https://www.humanconnectome.org). They have released behavioral data spanning myriad domains including: *health and family histo*ry (physical health assessments, menstrual cycle factors, family history of psychiatric and neurological disorders), *alertness* (Mini Mental Status Exam, Pittsburgh Sleep Questionnaire), *cognition* (episodic memory, executive function - cognitive flexibility and inhibitory processing, fluid intelligence, language – reading, decoding and vocabulary comprehension, processing speed, impulsivity, spatial orientation, sustained attention, verbal episodic memory, and working memory), *emotion* (emotion recognition, negative affect, psychological well-being, social relationships, stress and self efficacy), *motor function* (endurance, locomotion, dexterity, grip strength), *personality* (five factor NEO model), *psychiatric and life function* (Achenbach Adult Self-Report, syndrome scales and DSM-oriented scales, and psychiatric history), *sensory function* (audition, olfaction, pain, taste, vision, contrast sensitivity), and *substance use* (breathalyzer and drug test, alcohol and tobacco use 7-day retrospective, alcohol and tobacco use and dependence, illicit drug use, marijuana use and dependence). Twins and siblings were recruited for the study, with full genotyping data to be released soon (https://humanconnectome.org/about/pressroom/). A full catalog of the assessments done by the HCP that are available through the HCP ConnectomeDB database for housing and disseminating HCP data can be found in their online Data Dictionary (https://wiki.humanconnectome.org/display/PublicData/HCP+Data+Dictionary+Public-+500+Subject+Release) and in Hodge et al. (2016).^25^

Multi-modal neuroimaging data are also available from the HCP, including high spatial resolution structural and diffusion tensor imaging data and high spatial and temporal resolution task and resting state functional MRI (fMRI) data at 3T. 3T data were collected in over 1100 participants, retest data in 46 participants, multimodal 7T MRI data in 184 participants, and magnetoencephalography (MEG) data in 95 participants. Task fMRI data were collected during seven different tasks chosen to cover multiple domains of function and optimized to activate as many functional nodes, or regions of the brain, as possible.^26^ In addition, the HCP collected four 15 minute runs of r-fMRI data in each participant.^27^ Notably, the raw neuroimaging data and minimally pre-processed denoised resting state data, pre-processed and first-level statistical parametric maps from general linear modeling of the task fMRI data, and soon to be released genetics data are all available to researchers. HCP data are accessible by ordering the Connectome-In-A-Box (http://www.humanconnectome.org/data/connectome-in-a-box.html) directly from the HCP (to create local installations of the HCP data) or by accessing the data via the cloud through Amazon Web Services, which allows users to analyze HCP data without having to set up a local installation (which would require 100 TB of disk space). In addition, HCP data and analysis pipelines are detailed in several published reports^22,26,27,28^ and all imaging data processing scripts have been provided by the HCP to facilitate standardization of analysis methods. Details of the informatics tools developed by the HCP to enable high throughput data collection, automated analysis, and data sharing can be found in published reports.^25,29^

The ConnectomeDB, which is the platform for housing and disseminating the HCP data, is the foundation for the Connectome Coordination Facility (CCF). The CCF is an NIH-supported dissemination platform for the HCP data and data from numerous other NIH-funded HCP harmonized studies that are currently being conducted that will also be open access, including HCP Lifespan Babies, HCP Lifespan Children, HCP Aging, Developing HCP, as well as data from thirteen additional large-scale HCP-harmonized studies funded through the Connectomes of Human Diseases mechanism (https://grants.nih.gov/grants/guide/pa-files/PAR-14-281.html) covering diverse illnesses, such as Alzheimer’s Disease, epilepsy, frontotemporal degeneration, anxious misery, early psychosis, low vision and blindness, anxiety and depression. Once all of the HCP-style data from these studies are made available through the ConnectomeDB, it will be an amazing resource of data collected across studies that are all harmonized with each other, including overlapping deep-phenotyping and neuroimaging protocols, that can be used to replicate studies involving not only behaviors and the brain connectome in healthy individuals, but also in disease and development. Given the wide range of populations, diseases, and data from the HCP-style studies that will eventually be open access, raising awareness of the HCP itself for conducting replication studies will surely inspire graduate students, post-doctoral researchers, and investigators to consider replicating their own research and the research of others as an enhancement to their own work. In the rest of this research report, I demonstrate the power of the HCP imaging data for replication studies by using the neuroimaging data to replicate a seminal research report that has had a very high impact on the neuroimaging and neuroscience community but would be very challenging to replicate without access to data like the HCP.

In 2009, Smith et al published a paper in the Proceedings of the National Academy of Sciences^30^ that showed that the collection of brain networks that are “active” while a person is resting and engaged in idle thought (e.g., resting state networks or RSNs) correspond to the same functional networks used by the brain to perform tasks. This study showed for the first time the extent to which the set of RSNs consistently observable using fMRI during rest match the functional networks utilized by the brain during tasks and provided strong supporting evidence to an emerging literature showing that RSNs observable with fMRI were not simply due to non-neural physiological effects. Smith et al has been cited at least 1,550 times, in the top 1% of highly cited papers in Neuroscience and Behavior (Web of Science) and possibly more than 2,200 times (Google Scholar), underscoring its importance to the neuroscience and neuroimaging communities. Given the significance of this study, it would be reassuring if the main findings were replicated in an independent study. However, the Smith et al. study is challenging to replicate because one needs imaging data collected during the performance of many different tasks covering myriad behavioral domains to identify the full repertoire of task networks. Smith et al. were able to utilize the BrainMap database^31^ in combination with resting state fMRI data to investigate the links between RSNs and task activation networks. Using the BrainMap database, which is the largest database of human brain activation study results obtained using neuroimaging techniques, allowed them to pool together imaging results reported from more than 1600 published studies of brain function over a wide range of experimental paradigms to derive the full repertoire of the brain’s functional task networks. Thus, to replicate the findings in the Smith et al. study, one would need access to task fMRI data spanning many different tasks and functional domains, and resting state fMRI data. The HCP imaging data represent just such a dataset, with fMRI data collected during seven different tasks chosen to cover multiple domains of function, including emotion processing, incentive processing, language, motor, relational processing, social cognition, and working memory, and at rest, and are thus these data are ideally suited for replicating the Smith et al. study.

The Smith et al. study has not been particularly controversial because it is supported by converging evidence from many other studies and lines of research.^32,33,34^ However, replication of such an influential study is still critically important. For example, consider another highly cited influential study conducted by Strack, Martin & Stepper (1998) on the facial feedback hypothesis. That study has been cited more than 1600 times (Google Scholar) and is a common concept in introductory psychology texts and courses. This hypothesis is also supported by numerous related studies, and as such, became readily accepted in the psychology community. However, seventeen subsequent attempts to replicate the results resulted in heterogeneous findings, which motived a recent registered research report to conduct a meta-analysis of the replications.^35^ In the end, the meta-analysis was not able to replicate the findings of this seminal, widely accepted research study, underscoring the importance of replication of seminal influential studies, even if the findings are readily accepted based on the reputation of the scientist(s) who conducted the study or the journal in which the study is published, or on converging lines of evidence that support the conclusions of the study.

## Results

Smith et al. applied independent component analysis (ICA) to the BrainMap data and to resting state fMRI data to derive 20 independent components from each dataset representing both large-scale brain networks and artifact-related effects. He then identified task networks that corresponded to commonly observed RSNs^36,37^ using spatial cross-correlation (Smith et al. Figure 1). In the present study, these Smith et al. main findings were highly reproducible using the HCP task and resting state data. The spatial cross correlations between the HCP task networks and Smith’s BrainMap networks ranged from r = 0.26-0.74, with the minimum r=0.26 being highly significant (p=4×10^-4^, corrected). The spatial cross correlations between the HCP task and HCP RSNs ranged from 0.44-0.81, with the minimum r=0.44 being highly significant (p<4×10^-4^, corrected). The correspondence between the HCP task and HCP RSNs is much higher than the correspondence between the BrainMap networks and the RSNs reported in Smith et al. (r=0.26, p<1×10^-5^, corrected), although both are highly significant. This is due to the different nature of the task data that was used for each study (BrainMap pseudo-activation maps versus actual task activation Z-stat maps in the present study).

**Figure. 1.**
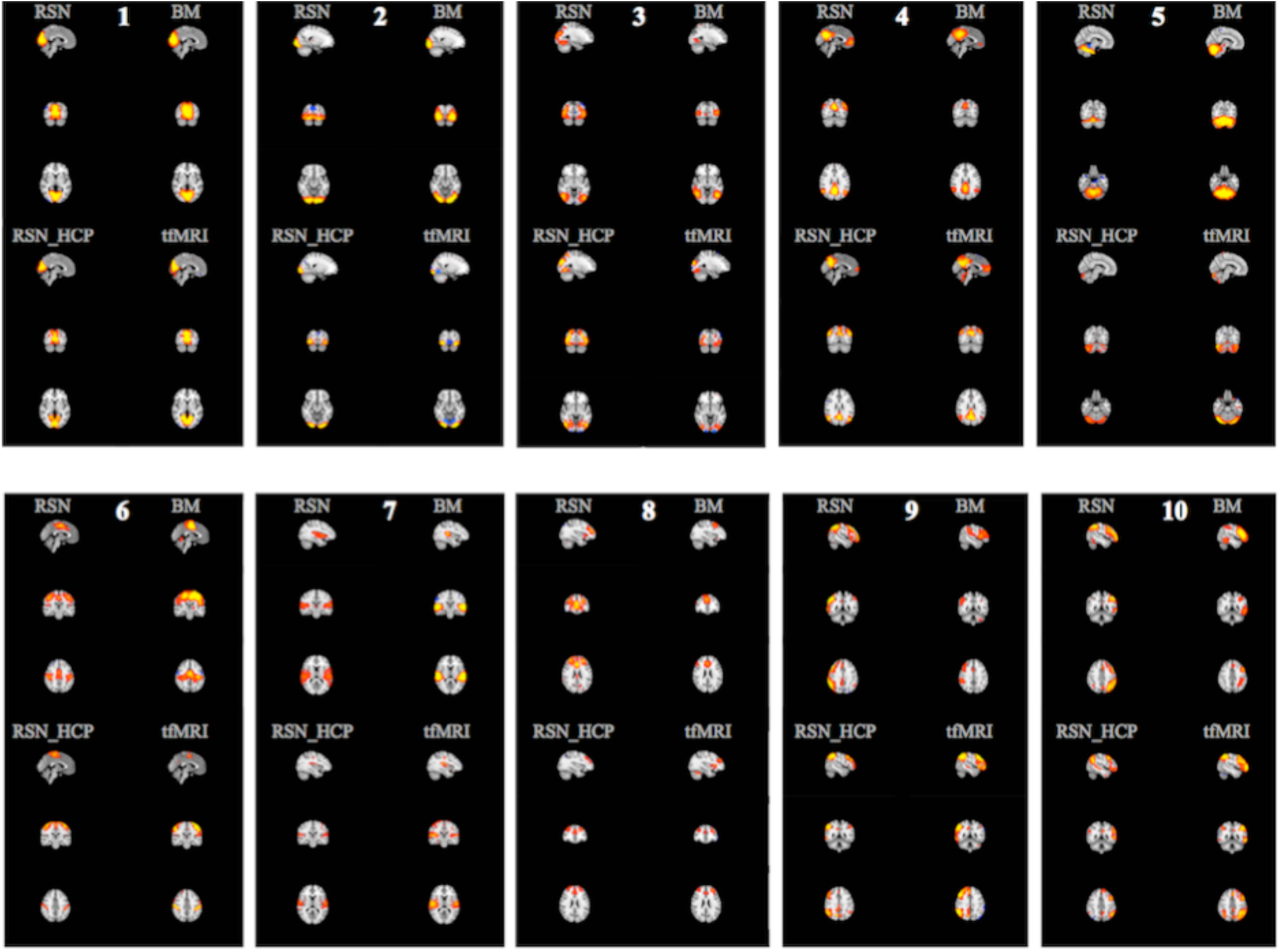
RSNs and BrainMap-derived task networks (top left and right of each panel, respectively) for ten networks reported in Smith et al. (2009) and corresponding RSNs and task networks derived using HCP data (bottom left and right of each panel, respectively). Networks are shown in red-yellow/blue-light blue, thresholded z>3. Networks are: 1) medial visual, 2) occipital pole, 3) lateral occipital, 4) default mode, 5) cerebellum, 6) sensorimotor, 7) auditory, 8) executive control, 9) right frontoparietal, 10) left frontoparietal.

Figure 1 shows all four sets of network maps, the HCP RSN and HCP task networks from the present study and the RSN and BrainMap networks from Smith et al. study (their Figure 1). The Smith et al. maps displayed alongside the HCP maps were constructed from their data files available at http://fsl.fmrib.ox.ac.uk/analysis/brainmap+rsns/. For Figure 1, all ICA Z-statistic maps were thresholded with Z=3.0 (the same as in Smith et al. Figure 1) and red-yellow and blue-light blue colors indicate the network (positive and negative values, respectively), overlaid onto the MNI standard brain image. The Z=3.0 threshold used in Smith et al. is based on an alternative hypothesis testing approach which applies a Gaussian-Gamma mixture model to the independent component spatial maps to determine the threshold for each map.^38^ In this case, a threshold of p=0.5 will achieve an equal probability of obtaining a false positive or a false negative (e.g., of a given voxel being in the background signal or the IC signal). Mixture modeling to threshold ICA maps is used to address the fact that spatial maps derived from the fixed-point iteration ICA algorithm in FSL MELODIC (and from Infomax or other similar algorithms) are optimized for maximal non-Gaussianity of the distribution of spatial intensities. In this case, simple transformation of ICA maps to Z scores and subsequent thresholding will not provide control of the false-positive rate. Using mixture modeling allows for such control and Z = 3 is approximately the average Z value that one obtains from thresholding a typical group ICA spatial map at p=0.5 when 20 components have been estimated (e.g., if more components are estimated, the threshold will increase due to reduced residuals). Applying a mixture model to the ICA maps, with p=0.5, in the present study gave an average value of Z ~ 2.5 so the threshold used to create the figure corresponds to a slightly greater probability of a voxel being signal rather than noise for the HCP-derived RSN and task network maps.

Correspondence between HCP and Smith et al. networks were similarly reproducible for the 70-component analysis, which results in a finer parcellation of the brain into sub-networks as compared with 20 estimated components (Smith et al. Figure 3; eight occipital and two sensorimotor networks). Spatial cross correlations between the RSNs from Smith et al. and the HCP RSNs ranged from 0.36 – 0.69 (results not shown).

It is not possible to reproduce how strongly each component in Figure 1 relates to the 66 possible behavioral domains as shown in Smith et al. (their Figure 2) due to the limited number of tasks in the present study. However, it is possible to determine the *magnitude* of network activation during each task component for the tasks in the HCP. In the BrainMap data, only the location in standard space of an activation blob is encoded in the BrainMap database with no spatial extent or magnitude information, however, many different tasks in any given behavioral domain are represented in the database. Smith et al. estimated a measure related to the frequency with which a spatial pattern was observed during related tasks in a given behavioral domain. While this metric is informative of which networks are engaged by tasks falling within a given domain, there is no information about how strong a network may be activated or suppressed during a particular task. In the present study, the strength of activation of each network for each task component, covering 29 different behavioral domains, was computed as follows. The set of spatial maps from the ICA of the HCP task Z-stat maps (contrast maps of brain activation during different task conditions) were used in a multiple spatial regression against each set of task Z-stat maps to extract a set of “subject courses” for that particular task component. These subject courses reflect the strength of the particular network activation in each subject, and can be averaged over all subjects for a particular task to compute the average magnitude of activation of each network during a task. This approach is different than the approach used to compute the matrix values shown in Smith et al. Figure 2. By using subject-specific task Z-stat maps and the multiple spatial regression approach in the present study, the magnitude of the activation of a network during a task component is preserved.

**Figure. 2:**
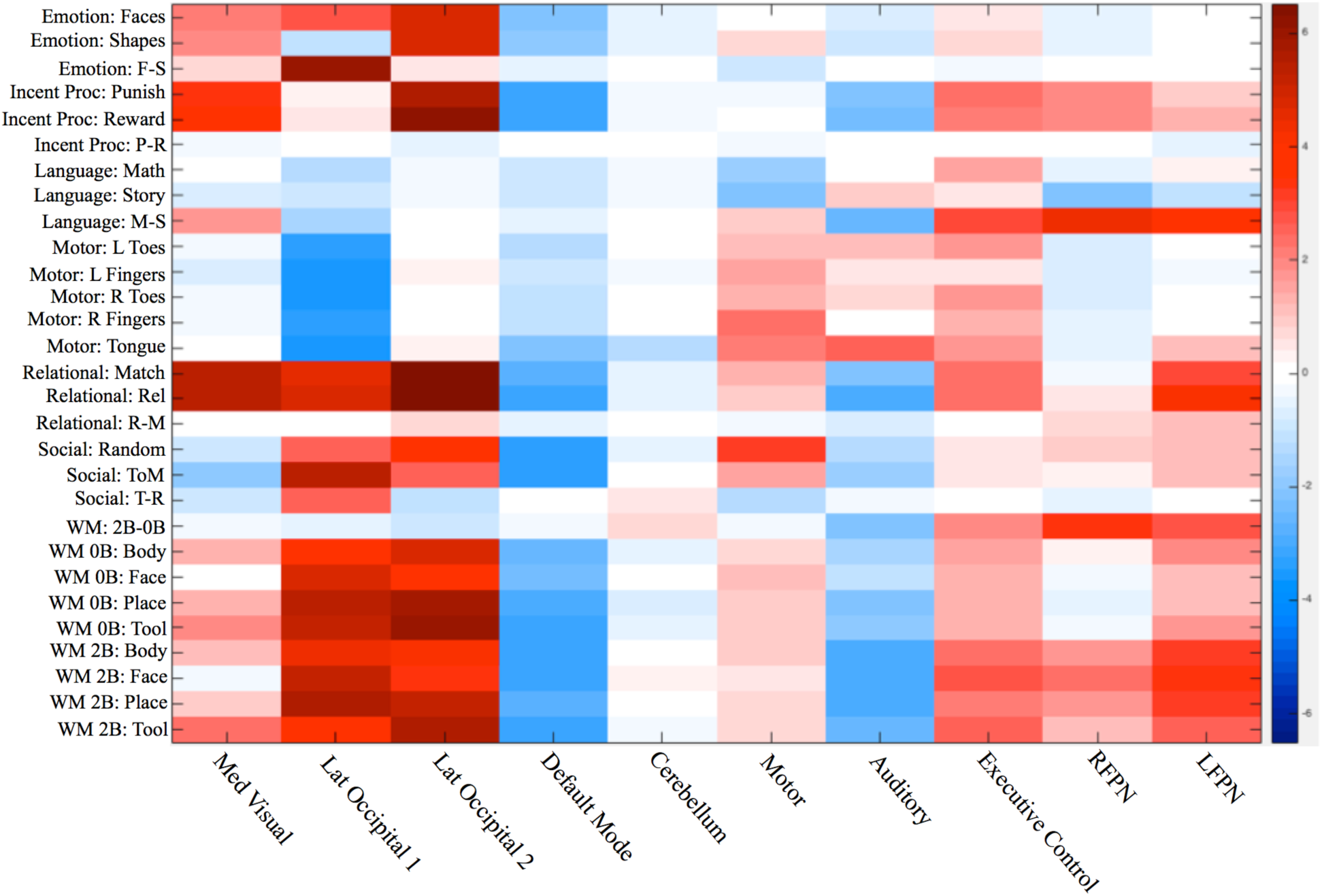
Magnitude ofactivation (Z-value) of each network during each task component.

Network activation magnitudes are displayed in Figure 2. No inference was done on this matrix as it is meant to parallel directly the qualitative results shown in Smith et al. Figure 2. However, some similarities with the Smith et al. Figure 2 are apparent. In Smith Figure 2, three neurocognitive networks, the executive control network and two lateralized left and right fronto-parietal networks, LFPN and RFPN, have high values for the domain Cognitive_Memory_Working. In my Figure 2, these three networks and cerebellum all showed a statistically significant activation during the working memory task (2 Back – 0 Back (2B-0B) condition), with p = 0.0001, taking into account family structure, corrected for number of networks and contrasts (e.g., 2-sided tests, with differences in reaction times and accuracy between 2B and 0B included as covariates of interest in the general linear model). All other networks showed statistically significant suppression during 2B-0B (p<0.02, corrected), which isn’t captured in the Smith matrix. As further validation, I also found that increasing network activation magnitude was associated with increases in reaction times during 2B relative to 0B for the executive control network, RFPN, and LFPN (p=0.0001, corrected) suggesting that as the working memory load increases, these networks increase their activity. The executive control network is comprised of dorsal anterior cingulate cortex (dACC), medial superior frontal cortex (msFC), and bilateral anterior insula/frontal operculum and has been shown to be a core system for the implementation of task sets that provides stable “set-maintenance” over entire task epochs over a variety of tasks.^39,40^ In the present study, this network shows increased activity during nearly all of the tasks (relative to their lower level control conditions), which is consistent with being a core system as described by Dosenbach et al. (2006) and with Smith’s findings that this network corresponded to several cognition paradigms. The LFPN and RFPN are posited by Dosenbach et al. (2008) to be control networks (e.g., a single network in their study, but split into two lateralized networks in our study and in other ICA-based fMRI studies) that potentially initiate and adjust control on a trial-to-trial basis and respond to events that carry performance feedback information. The prefrontal and parietal regions in all three of these networks have been implicated in previous studies of working memory^41^ and both networks were reported to show preference for n-back working memory tasks in further work (co-authored by Dr. Smith) that built upon the Smith et al study.^42^

Figure 2 results also shows that the default mode network (DMN) is suppressed during most tasks, which was also observed by Smith et al (stated in the text, not captured in his Figure 2). There appears to be stronger suppression of the DMN for those tasks with presumably greater cognitive load, which is consistent with several studies showing that the magnitude of activity within the DMN during task performance is related to task-load during brain activation.^43,44^ Occipital networks are activated during tasks in which there are visual stimuli, but not when the tasks involved auditory stimuli, e.g., in the math and story blocks of the language task, or the motor tasks, which is also consistent with the Smith et al. findings in his Figure 2.

There is a subtle point about the results shown in the present Figure 2. Namely, all of the networks that are activated (or suppressed) during a particular task are identified using the regression-based analysis approach used in the present study, and the activity of a specific task network during each task component is determined with all other network activity “partialled out” by virtue of the multiple spatial regression of all network maps against the subject activation maps. This means that the activity in each network is separately assessed – and that the specific set of networks that are engaged during a given task can be determined. This is in contrast to activation maps from a standard voxel-wise general linear model analysis, which instead show all of the brain regions that are activated during the task, aggregated together into a single map.

Thus, the resulting map from a whole-brain voxelwise GLM analysis reflects a composite of regions that constitute different networks, without differentiating each network itself. For example, consider again the working memory (2B-0B) contrast. In Barch et al. (2013), the group activation map obtained using a multi-level general linear model shows deactivations in medial prefrontal cortex and auditory regions (Figure 3 in Barch et al.). It is unknown whether the activated regions represent one or more networks in the aggregate activation map that results from the voxel-wise GLM analysis. Using the analysis approach in the present study, network activity can be disentangled, and we conclude that executive control, LFPN, and RFPN are all activated during 2B-0B. The auditory network is also strongly suppressed during this condition (and during many other conditions, possibly due to a need to block out distracting scanner noise). Thus, using the analysis approach utilized in the present study allows for each network that may be activated (or suppressed) by the task to be studied separately from other networks.

## Discussion

Replication studies play a key role in efficient science by testing directly the stability of novel findings in a rigorous scientific way, that is less influenced by factors (such as publication bias) that can propagate spurious results or practices into a field. One of the aims of the present study was to bring attention to the HCP as a goldmine of behavioral data, neuroimaging data, and (soon) genetic data that can be used for replication studies. Importantly, data are being collected in numerous ongoing NIH-funded HCP-harmonized studies of the lifespan, development, and diseases that also will be made available to researchers through the HCP informatics infrastructure as the studies complete, thus collectively providing extensive opportunities for replication studies across diverse human research domains. Hopefully, knowledge of the depth and breadth of the HCP data will inspire investigators to think about ways these open data can be used to support their research endeavors. In this report, I demonstrated how the diversity of the fMRI data in the HCP, covering seven different tasks and resting state, could be used to replicate the complex multi-modal Smith et al. (2009) neuroimaging study. Reassuringly, the Smith et al. findings were replicable under the most challenging form of generalizability, using different subjects, stimuli, and analysis methods,^15^ which moves the field forward by providing additional critical support to the findings by Smith et al. My findings are in agreement with the Smith et al. interpretation of RSNs measured using fMRI as reflecting neural activity, not just hemodynamics and other physiological processes, and as corresponding to task activation networks. While the correspondence between resting state and task brain networks is now generally accepted within the neuroimaging community, it is always good to revisit studies that may involve truly novel analysis methods and/or significant findings that shape the field.

## Methods

### I. Human Connectome Project

The second major release of the HCP data collected in 500 healthy adults (aged 22-35) was used for the current study. Individuals with severe neurodevelopmental disorders, neuropsychiatric disorders, or neurologic disorders, or with illnesses such as diabetes and high blood pressure, were excluded from the HCP study. MRI scanning was done using a customized 3T Siemens Connectome Skyra using a standard 32-channel Siemens receive head coil and a body transmission coil. T1-weighted high resolution structural images acquired using a 3D MPRAGE sequence with 0.7 mm isotropic resolution (FOV = 224 mm, matrix = 320, 256 sagittal slices, TR = 2400 ms, TE = 2.14 ms, TI = 1000 ms, FA = 8°) were used in the HCP minimal pre-processing pipelines to register functional MRI data to a standard brain space. Resting state fMRI data were collected using gradient-echo echo-planar imaging (EPI) with 2.0 mm isotropic resolution (FOV=208x180 mm, matrix =104x90, 72 slices, TR = 720 ms, TE = 33.1 ms, FA = 52°, multi-band factor = 8, 1200 frames, ~15 min/run). Task fMRI data were collected using the same scanning sequence as the resting state fMRI data, although the number of frames per run (with 2 runs/task) varied from task to task. Runs with left-right and right-left phase encoding were done for both resting state and task fMRI to correct for EPI distortions. All subject recruitment procedures and informed consent forms, including consent to share de-identified data, were approved by the Washington University Institutional Review Board (IRB). See Glasser et al., 2016. For the present study, after permission was obtained from the HCP to use the Open Access and Restricted Access data for the present study (see Data Availability Statement below), a protocol filed with the McLean Hospital Institutional Review Board (IRB) met criteria for exemption.

### II. Identification of Task Networks from HCP Task fMRI Data

Task fMRI data were utilized from seven different tasks: emotion processing, incentive processing/gambling, language, motor, relational processing, social cognition, and working memory. I used volumetric outcomes from the minimal pre-processing pipelines developed by the HCP^45^. For the HCP minimal pre-processing pipeline, the task fMRI data for each subject underwent corrections for gradient distortions, subject motion, and echo-planar imaging (EPI) distortions, and were also registered to the subject’s high-resolution T1-weighted MRI. All corrections and the transformation of the fMRI data to MNI standard space (via non-linear transformation of the subject’s T1-weighted structural MRI into MNI standard space) were implemented in a single resampling step using the transforms for each registration step (fMRI to T1 and T1 to MNI) and the distortion corrections.

First-level statistical modeling was also implemented by the HCP. The pre-processed fMRI timeseries at each voxel (or spatial location) in the task fMRI data was fit with a general linear model (GLM). Regressors that modeled the brain’s fMRI signal in response to the task conditions were included in the model. 3D spatial maps (e.g., one value per voxel in the brain) of contrasts of the parameter estimates (COPEs) were computed corresponding to the average activation during each task component and to differences in activation between different task components (e.g., subtraction images). Notably, these COPE maps reflect the magnitude of the brain activation between two task conditions.

Twenty-five COPEs were selected from across tasks to be fed into a group independent component analysis (ICA) to identify task networks. Table 1 describes each task COPE, with 3-5 COPEs per task.

**Table 1.**

| Task | HCP Cope # | Behavioral Domain |
| --- | --- | --- |
| **Emotion Processing** | 1 | *Angry/Fearful Faces* (which of two faces at the bottom of the screen match face at top of screen; faces are angry or fearful) |
|  | 2 | *Shapes* (which of two shapes at the bottom of the screen match shape at top of screen) |
|  | 3 | *Angry/Fearful Faces – Shapes* |
| **Incentive Processing/Gambling** | 1 | *Punishment* (lose money when guessing the number on a mystery card) |
|  | 2 | *Reward* (win money when guessing the number on a mystery card) |
|  | 3 | *Punishment – Reward* |
| **Language** | 1 | *Math Problems* (auditory presentation of math problems, requires subjects to complete addition/subtraction problems) |
|  | 2 | *Stories* (auditory presentation of stories, participants answer questions about the story) |
|  | 3 | *Math Problems – Stories* |
| **Motor** | 2 | *Squeeze Left Toes* |
|  | 3 | *Tap Left Fingers* |
|  | 4 | *Squeeze Right Toes* |
|  | 5 | *Tap Right Fingers* |
|  | 6 | *Move Tongue* |
| **Relational Processing** | 1 | *Match Condition* (does single object at bottom of screen match either of two objects at top of screen on either shape or texture?) |
|  | 2 | *Relational Condition* (does bottom pair of objects vary on the same dimension (shape or texture) as top pair of objects?) |
|  | 4 | *Relational – Match* |
| **Social Cognition (Theory of Mind)** | 1 | *Random* (participants judge video clips of objects to be either interacting in some way or moving randomly; in this condition, objects are moving randomly) |
|  | 2 | *Theory of Mind: ToM* (participants judge video clips of objects to be either interacting in some way or moving randomly; in this condition, objects are interacting) |
|  | 6 | *ToM – Random* |
| **Working Memory\*** | 11 | *2 Back – 0 Back* (blocks of trials consisting of pictures, faces, places, tools, and body parts, with ½ of blocks using 2-back working memory task and ½ blocks using 0-back working memory task being contrasted) |
|  | 5 | *Body Parts (0 Back blocks) vs. Baseline* |
|  | 6 | *Faces (0 Back blocks) vs. Baseline* |
|  | 7 | *Places (0 Back blocks) vs. Baseline* |
|  | 8 | *Tools (0 Back blocks) vs. Baseline* |
|  | 1 | *Body Parts (2 Back blocks) vs. Baseline* |
|  | 2 | *Faces (2 Back blocks) vs. Baseline* |
|  | 3 | *Places (2 Back blocks) vs. Baseline* |
|  | 4 | *Tools (2 Back blocks) vs. Baseline* |
| *For the ICA, working memory contrasts that averaged 0 Back and 2 Back blocks together for each stimulus type were used (COPEs 15-18), but the COPEs for 0 Back and 2 Back blocks relative to baseline were used to analyze network activation to facilitate interpretability. |  |  |

The sample size for each task is as follows:

- Emotion Processing. 452 subjects, 3 COPES, 1,356 total images
- Incentive Processing. 449 subjects, 3 COPES, 1,347 total images
- Language. 433 subjects, 3 COPES, 1,299 total images
- Motor. 415 subjects, 5 COPES, 2,075 total images
- Relational Processing. 435 subjects, 3 COPES, 1,305 total images
- Social Cognition. 452 subjects, 3 COPES, 1,356 total images
- Working Memory. 411 subjects, 5 COPES, 2,055 total images

The number of subjects varies for each task because participants with greater than 2 mm of motion (maximum absolute root mean square) in any task run led to exclusion of their COPE map from the analysis. E.g., two runs were done for each task (with different phase encoding directions to correct for EPI distortions), with average COPE maps being calculated using a second-level GLM. These average COPE maps are used in the present analysis.

All 10,793 COPE maps were fed into a group ICA conducted using FSL MELODIC^38^. Two group ICAs were conducted, one with twenty and one with seventy components estimated from the group ICA, the same as in the Smith et al. study, to identify large-scale brain networks and to do a finer parcellation, respectively. The spatial independent component maps were thresholded using a Gaussian-Gamma mixture model with p=0.5 such that equal weight was given to obtaining either a false positive or a false negative in the spatial map. Note that Smith et al., constructed 7,342 activation-peak images (pseudo-brain activation maps constructed by filling an empty image with points corresponding to reported standard space coordinates of statistically significant local maxima in the activation maps from the original study that are archived in the BrainMap database, then convolving these points with a Gaussian kernel to mimic spatial extent), which were submitted to a group ICA. Thus, the number of maps used in the present study is the same order of magnitude as the number of maps used in the Smith et al. study.

Task activation networks were identified corresponding to those from Smith et al. (e.g., from the ICA of the BrainMap data) by visual inspection and spatial cross-correlation. Significance of cross-correlations was determined as follows. The corrected p-value was computed based on a Bonferroni correction for the number of possible paired comparisons (400 for HCP task vs Smith task; 400 for HCP task vs HCP rest) and a correction for the spatial degrees of freedom using Gaussian random field theory and an empirical smoothness estimation (average number of resels = 322 for HCP task spatial maps, which was lower than both the average for the BrainMap task maps (2143 resels) and the average for the HCP RSNs (375)). For example, the correlation probability for r = 0.26 with 322 degrees of freedom is p=1×10^-6^ (one-sided), multiplying by this value by 400 gives p=4×10^-4^ corrected. See Smith et al. for more details.

The activation magnitudes for each network during each task component were computed as follows. Multiple spatial regression of the HCP dimensionality 20 ICA maps (e.g., all 20 maps together) against the 4D file of COPE maps, e.g., with one COPE map for each subject concatenated across all N subjects, was done for each task component. For each COPE file, the resulting regression parameters are a “subject-series” of loadings (one per subject) that were averaged together to give the average network activity during that component of the task. The regression parameters are Z-statistics since the COPE Z-stat maps from the HCP first-level analysis were used for the analysis. The values in Figure 2 were computed as the average of the subject-series of loadings for each COPE. To assess the activated networks during the 2 Back (2B) vs 0 Back (0B) contrast and the relationships with change in accuracy (0B-2B) and change in reaction time (2B-0B), a general linear model was implemented with the subject-series for each network for the 2B-0B as dependent variables and change in %accuracy and change in reaction times as covariates using PALM (Permutation Analysis of Linear Models),^46^ which also takes into account the family structure of the HCP data, and to correct for the number of networks and contrasts. Exchangeability blocks were determined that captured family structure to determine acceptable permutations, and 10,000 permutations were done.

### III. Identification of Resting State Networks from HCP Resting State fMRI Data

RSNs were identified from the HCP data using outcomes from the minimal pre-processing pipeline of the resting state fMRI data that were provided by the HCP.^27^ Minimal pre-processing of resting state fMRI data included corrections for spatial distortions caused by gradient nonlinearities, head motion, B_0_ distortion, denoising using FSL FIX^47^ and registration to the T1-weighted structural image. All transforms were concatenated together with the T1 to MNI standard space transformation and applied to the resting state fMRI data in a single resampling step to register the corrected fMRI data to MNI standard space. fMRI data were also temporally filtered with a high pass filter and then each subject’s fMRI data was analyzed using spatial ICA, with the MELODIC algorithm estimating the number of components. In the HCP framework, these ICA maps are used to denoise the fMRI data prior to any subsequent resting state analyses. In the present analysis, these single-subject ICA maps, from 20 participants, were fed together into a group ICA using FSL MELODIC to identify the collection of RSNs that were common to the group of subjects. The single-subject ICA maps were used for the group ICA instead of the original minimally-preprocessed resting state fMRI data simply to reduce computational load, which is much greater with HCP data due to the extremely high spatial and temporal resolution of the fMRI data (2×2×2 mm^3^ with 0.75 second sampling intervals, 15 minutes/run, 1200 volumes/run). As an aside, this is the same order of magnitude of participants in the original study by Smith et al., which utilized resting state fMRI data from 36 participants.

Two ICAs were done, with the number of components fixed to twenty and seventy (as in Smith et al.) and RSNs that corresponded to the RSNs in Smith et al. were identified by visual inspection and spatial cross-correlation. For the 20-component ICA, the spatial cross-correlations between the HCP RSNs and the Smith RSNs (for 10 networks shown in Smith Figure 1) ranged from 0.43-0.74 (except for the cerebellum network which was cut off in Smith’s data, but fully covered in the HCP data, resulting in a spatial cross correlation of 0.33). The spatial cross-correlations between HCP RSNs and HCP task networks (Figure 1) ranged from 0.44-0.81. The minimum spatial correlation of 0.44 is even greater than the minimum reported in Smith et al., r=0.25, and so is even more highly significant than p<5×10^-6^ (although I did not estimate the actual p value since it’s not necessary given the high level of significance).

### IV. Data Availability

All data used in the present study are available for download from the Human Connectome Project (www.humanconnectome.org). Users must agree to data use terms for the HCP before being allowed access to the data and ConnectomeDB, details are provided at https://www.humanconnectome.org/study/hcp-young-adult/data-use-terms.

The HCP has implemented a two-tiered plan for data sharing, with different provisions for handling Open Access data and Restricted data (e.g., data related to family structure, age by year, handedness, etc). Both Open Access and Restricted data were utilized in the present study. The resting state and task fMRI outcomes provided from the HCP processing pipelines and the in-scanner task performance measures for the working memory task are Open Access, thus users must be granted first-tier permission by the HCP to access that data. However, the family structure information that was utilized in the present study to do inference with non-parametric permutation methods described in Sections II and III is Restricted data, which would require second-tier permission by the HCP to access that information. In addition, the HCP requires that users “must abide by a prohibition against publishing certain types of individual data in combination that could make an individual recognizable or that could harm and embarrass someone who was inadvertently identified” as per the Restricted Data Use Terms and application. See https://www.humanconnectome.org/study/hcp-young-adult/document/restricted-data-usage for more details.

Users must also consult with their local IRB or Ethics Committee (EC) before utilizing the HCP data to ensure that IRB or EC approval is not needed before beginning research with the HCP data. If needed, and upon request, the HCP will provide a certificate to users confirming acceptance of the HCP Open and Restricted Access Data Use Terms. See https://www.humanconnectome.org/study/hcp-young-adult/data-use-terms.

## V. Code Availability

Matlab code to produce results (for Figure 2 and working memory) is available upon request.

## Acknowledgements

This work was supported by the National Institutes of Health (LDN: DA037265, AA024565).

Data were provided by the Human Connectome Project WU-Minn Consortium (Principal Investigators: David Van Essen and Kamil Ugurbil, 1U54MH091657). The Human Connectome Project was supported by the 16 NIH Institutes and Centers that support the NIH Blueprint for Neuroscience Research; and by the McDonnell Center for Systems Neuroscience at Washington University.

